# Interferon-gamma mediates skeletal muscle lesions through JAK/STAT pathway activation in inclusion body myositis

**DOI:** 10.1101/2021.12.16.472927

**Authors:** Cyrielle Hou, Baptiste Periou, Marianne Gervais, Juliette Berthier, Yasmine Baba-Amer, Sarah Souvannanorath, Edoardo Malfatti, Fréderic Relaix, Maximilien Bencze, François Jérôme Authier

**Affiliations:** Univ Paris Est Créteil, INSERM, IMRB, F-94010 Créteil, France; Reference Centre for Neuromuscular Diseases “Nord-Est-Ile de France”, FILNEMUS, France; AP-HP, Hôpital Mondor, Service d’histologie, F-94010 Créteil, France

**Keywords:** Inclusion body myositis (IBM), Inflammatory myopathies, IFNγ, JAK-STAT, skeletal muscle, myogenesis, satellite cells

## Abstract

Dysimmune and Inflammatory Myopathies (DIMs) are acquired idiopathic myopathy associated with immune response dysregulation. Inclusion Body Myositis (IBM), the most common DIMs, is characterized by endomysial infiltrates of cytotoxic T lymphocytes CD8, muscle type II-interferon (IFNγ) signature, and by the lack of response to immunomodulatory therapies. We showed that IBM was pathologically characterized by the presence of chronic degenerative myopathic features including myofiber atrophy, fibrosis, adipose involution, and the altered functions of skeletal muscle stem cells. Here, we demonstrated that protracted systemic exposure to IFNγ delayed muscle regeneration and led to IBM-like muscular degenerative changes in mice. *In vitro*, IFNγ treatment inhibited the activation, proliferation, migration, differentiation, and fusion of myogenic progenitor cells and promoted their senescence through JAK-STAT-dependent activation. Finally, JAK-STAT inhibitor, ruxolitinib abrogated the deleterious effects of IFNγ on muscle regeneration, suggesting that the JAK-STAT pathway could represent a new therapeutic target for IBM.

## Introduction

Dysimmune and Inflammatory Myopathies (DIMs) differ from each other by the profile of muscle tissue injuries and pathological mechanisms. Among DIMs, inclusion body myositis (IBM) is characterized by a slowly progressive muscle involvement with unique clinical and pathological features. IBM disease typically occurs over 50 years, with insidious progression and asymmetrical, proximodistal, muscle involvement mainly affecting finger flexors and quadriceps^1^. Unlike other DIMs, IBM is regarded as refractory to immunosuppressive therapies, and therefore probably one of the most disabling^2^. From a pathological view, IBM combines immune-mediated polymyositis-type inflammatory process with degenerative features including myofiber atrophy, amyloid deposits (β- APP), rimmed vacuoles, and fibrosis in the muscle^3,4^. Inflammatory infiltrates contain predominantly cytotoxic CD8+ T lymphocytes that mediate myonecrosis but the mechanisms underlying muscle dysfunction in IBM remains largely unknown^5^. In the vicinity of these infiltrates, muscle fibers in IBM abnormally express Major-Histocompatibility Class I (MHC-I) and class II (MHC-II) molecules on their surface^6^ that is associated with muscular type 2 IFN (IFNγ) signature^7–9^.

IFNγ is a potent inducer of MHC-II expression through the activation of JAK-STAT pathway and CIITA transactivator^10–12^. IFNγ is a cytokine that is produced by immune cells, including T lymphocytes and natural killer (NK) cells, and which is required for innate and adaptive immunity against infection. IFNγ is a mediator for macrophage polarization, lymphocyte regulation, proliferation, and survival^12,13^. On the other hand, the implication of IFNγ in muscle homeostasis and repair remains controversial. In mice, transient IFNγ injection improves muscle function and decreases fibrosis after laceration injury^14^. Inactivation of IFNγ *in vivo* negatively impacts muscle regeneration by impairing macrophage function, decreasing myogenic cell proliferation, and increasing fibrosis^15^. In addition, the overexpression of IFNγ at the neuromuscular junction induces necrotizing myopathy^16^ and the loss of IFNγ in mdx mice ameliorates their dystrophic phenotype. Finally, these data suggest a possible role for aberrant IFNγ signature in IBM-associated muscle damage.

To address this question, we examined transcriptomic and histological profiles of muscle biopsies from DIMs patients and analyzed *in vivo* and *in vitro* the impact of sustained IFNγ exposure on skeletal muscle tissue and myogenic cell behavior. We confirmed that IBM displays the strongest muscular IFNγ signature among DIMs. Experimentally, increased plasma IFNγ delayed post-injury muscle regeneration and induced IBM-like muscle changes including positive MHC-II myofibers, fibrosis, adipose involution, and myofiber atrophy. The deleterious effects of IFNγ were prevented by ruxolitinib, an inhibitor of the JAK-STAT pathway. *In vitro* IFNγ exposure promoted senescence and reduced activation, proliferation, migration, differentiation, and fusion of human muscle progenitor cells (MPC), all these effects being reversed by ruxolitinib treatment. Altogether, our data demonstrate that IBM muscles are characterized by an upregulation of the IFNγ signaling and that aberrant muscle IFNγ expression recapitulates IBM muscle phenotype, which can be reverted by targeting the JAK-STAT pathway. This work offers new therapeutic approaches for IBM patients using JAK-STAT inhibitors.

## Results

### IBM differs from other DIMs by its highest muscular IFNγ signature

We performed RNA-sequencing (RNA-seq) and histological study in muscle samples from healthy human controls (CTL, n=5), and patients with dermatomyositis (DM, n=5), anti-synthetase syndrome (ASS, n=5), and IBM (n=4). Gene expression profile in IBM strikingly differed from DM with 1816 genes differentially expressed, while only 190 genes differed between DM and ASS, and 14 genes between IBM and ASS *(Figure S1A)*. T-lymphocytes infiltrates are predominant in IBM muscle^18,19^. Gene ontology and analysis showed upregulated genes involved in T lymphocytes activation/proliferation, macrophage activation, and MHC-II protein expression in IBM muscles *(Figure 1A, B)*. IFN-I signaling was upregulated in DM muscles compared to the CTL, ASS, and IBM *(Figure S1B)*. In contrast, IFNγ expression and its IFNγ gene signaling were significantly upregulated in IBM and ASS, as confirmed by transcriptomic and RT-qPCR analysis *(Figure 1B, C)*. Interestingly, IFN-II-related genes were strongly overexpressed in IBM patients *(Figure 1B)*. DM presented some upregulated IFN-II-related genes compared to the CTL *(Figure 1B)* but the IFN-II signature was weaker in DM than in ASS and IBM patients. The expression of the *Hla-dr*, -dm, -dq genes was higher in IBM, moderately increased in ASS, and minimally increased in DM compared to CTL muscle biopsies *(Figure 1B, C)*. Overall, the level of regulation of *Mhc-II* expression is correlated to *Ifnγ* expression, especially in ASS and IBM patients *(Figure 1D)*.

**Figure 1.**
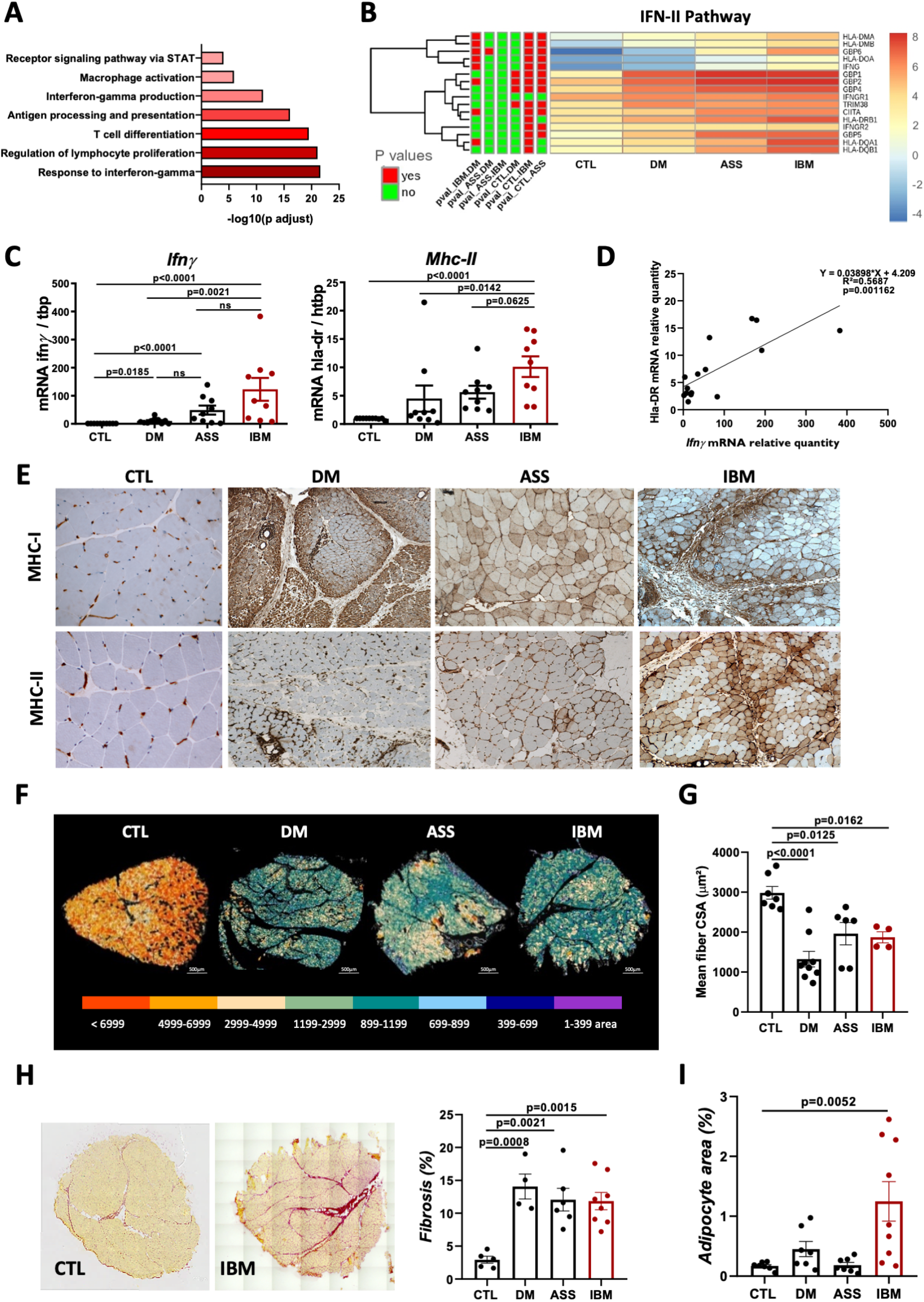
IBM differs from other DIMs by its highest muscular IFNγ signature. **(A)** Gene ontology analysis for biological processes of the upregulated (red) genes in Inclusion body myositis (IBM) muscle compared to control (CTL) muscles. Selected enriched terms are presented according to the fold enrichment. **(B)** Heatmap pathway IFN-II with the normalized reads per gene of CTL (n=5), Dermatomyositis (DM, n=5), Anti-synthetase syndrome (ASS, n=5) and IBM (n=4) RNA-seq data. **(C)** Quantification of *Ifnγ* and *Hla-Dr (Mhc-II)* genes expression by RT-qPCR between DM (n=10), ASS (n=9), IBM (n=9) and CTL (n=10) muscles. Mann-Whitney U test, means ± SEM. **(D)** Positive correlation between *Hla-Dr* and *Ifnγ* gene expressions in IBM and ASS muscles. Pearson’s correlation method. **(E)** Representative immunohistochemistry images showing MHC-I (top panel), and MHC-II staining (bottom panel) in IBM, ASS, DM and CTL muscles of our patient cohort with peroxidase (HRP)-conjugated polyclonal antibody). **(F)** Myofiber size analyses based on laminin immunostaining and automated analysis, as previously described^55^. **(G)** Cross-section-area (CSA) mean quantification in CTL (n=7), DM (n=9), ASS (n=6) and IBM (n=4) muscles. These myopathies are defined by myofibers atrophy, perfascicular in DM, and general in IBM. Means ± SEM, One-way ANOVA, Tukey’s multiple comparison. **(H)** Representative images of collagen deposit by Sirius Red staining (*left*) and quantification of the percentage of connective tissue area (*right*) performed CTL (n=5), DM (n=4), ASS (n=6) and IBM (n=8) muscles. Means ± SEM, One-way ANOVA, Tukey’s multiple comparison. **(I)** Quantification of adipocytes invasion performed on muscle sections from CTL (n=7), DM (n=7), ASS (n=7) and IBM patient (n=10). Means ± SEM. Mann-Whitney U test.

IFN-I/II cytokines are both potent inducers of MHC-I while IFNγ solely induces MHC-II expression through CIITA activation^10^. To histologically discriminate the type of IFN signature on CTL, DM, ASS, and IBM muscles, we performed immunostaining of MHC-I and MHC-II. Healthy CTL muscles, myofibers did not express MHC-I nor MHC-II proteins. In contrast, while all myofibers expressed MHC-I but not MHC-II in DM, myofibers expressed both MHC-I and MHC-II in ASS and IBM *(Figure 1E)*. Such MHC pattern confirmed that myofibers are mainly exposed to IFN-I in DM, and IFN-II in IBM and ASS patients. As assessed by 3D cleared muscle imaging, MHC-II labeling colocalized with dystrophin at the plasma membrane in IBM (*Figure S2A, supplemental videos*). This indicates that MHC-II is expressed at the sarcolemma of fully differentiated myofibers, making them potential targets for cytotoxic immune response^20^. We next looked at whether different IFN-I or -II signatures in DIMs patients were associated with specific histological muscle features. Regardless of the type of IFN signature, DIMs showed strong myofiber atrophy, as assessed by the decrease in myofiber size and fibrosis in DM, ASS and IBM *(Figure 1F-I)*. However, only IBM muscles displayed adipocytes muscle invasion *(Figure 1J)*.

### Systemic elevation of IFNγ induces IBM-like features in regenerating wild-type muscles

IFNγ is a labile cytokine with a short half-life *in vivo*^21,22^. To investigate the link between IFNγ and IBM pathophysiology, we implanted a subcutaneous osmotic pump releasing continuous recombinant mouse IFNγ in wild-type mice for 14 days. Prior to IFNγ delivery, *Tibialis Anterior* (TA) muscles were injured by BaCl_2_ injection^23^. Control injured animals received osmotic pumps releasing saline solution (CTL mice) *(Figure 2A)*. The efficacy of the systemic release of IFNγ in mice was confirmed by ELISA analysis. Two weeks following mouse surgery, mice implanted with IFNγ-containing pumps showed a 5-fold increase of IFNγ compared to saline-containing CTL mice *(Figure 2B)*. The IFNγ concentration reached in IFNγ mice was clinically relevant since it is comparable to the serum IFNγ level obtained in patients with infectious disease^24^. Increased systemic levels of IFNγ in mice were associated with a significant increase in muscle *CIIta* and *mhc-II* transcripts *(Figure 2C, D)* and MHC-II protein expression in the muscle at 7 days *(data not shown)* and 14 days post-injury *(Figure 2C, D)*. At 14 days, MHC-II protein expression was localized at the sarcolemma in IFNγ mice, as in IBM patients *(Figure S2A)*. IBM is associated with macrophage inflammatory infiltrates into the muscle. Since IFNγ promotes macrophage polarization and activates pro-inflammatory phenotype (M1)^12^, we thus explored the extent of macrophage infiltrates in IFNγ mice. Higher CD68+ macrophage density was observed in muscle from IFNγ mice. We found an increased expression of *iNos*, and *TGFß* expression in IFNγ mice muscles (*Figure 2F, G*), both transcripts being characteristic of M1 and M2 polarization, respectively. Macrophages and more specifically M2 subtypes are a strong producer of TGFβ, which is involved in muscle fibrosis^25^. Indeed, fibrosis was increased in IFNγ mice compared to CTL mice *(Figure 2H-K)*. Hence, the muscular content of adipocytes was increased in IFNγ mice compared to CTL mice at 14 days post-injury *(Figure 2J)*.

**Figure 2.**
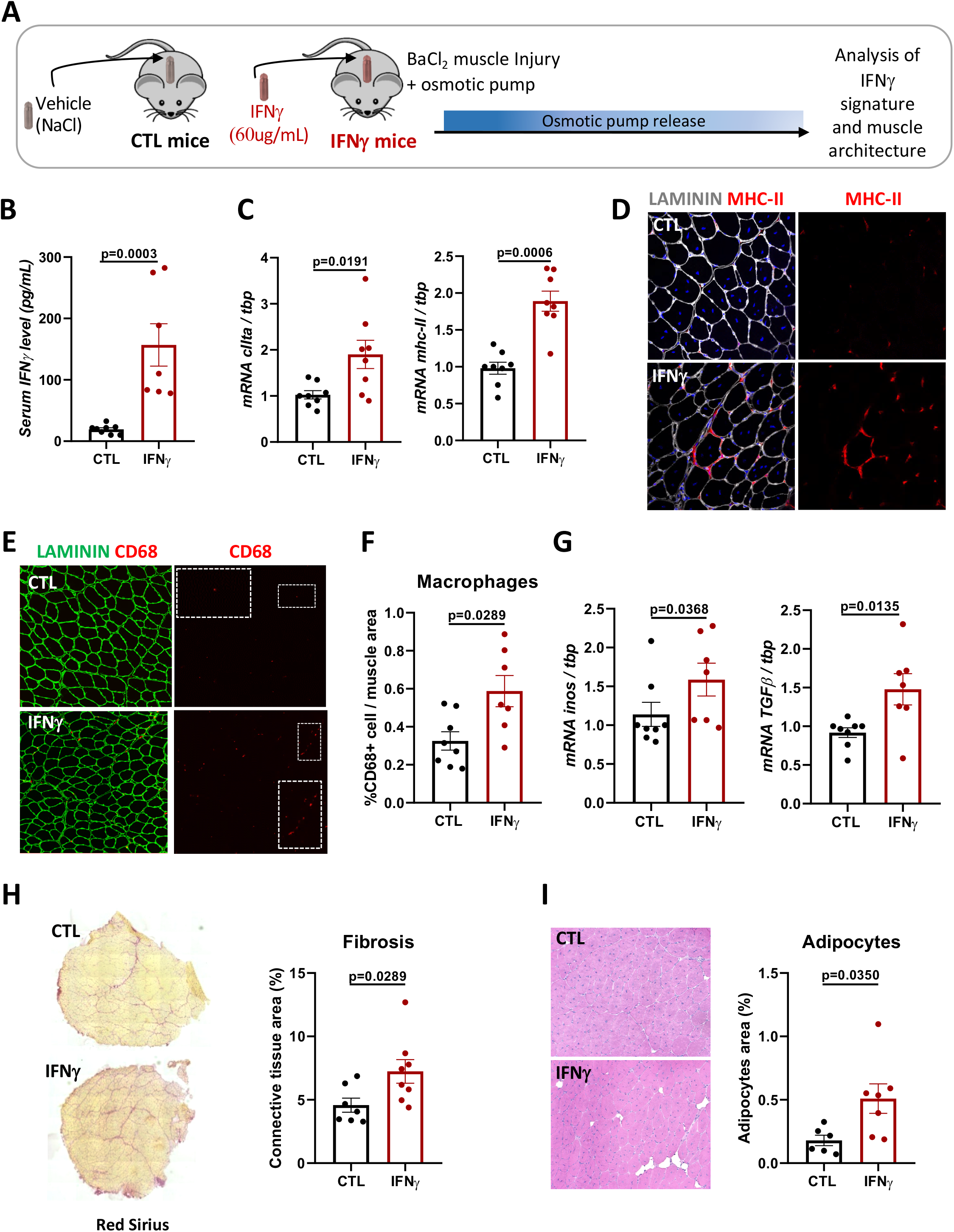
Systemic elevation of IFNγ delays myofiber regeneration in mice. **(A)** Experimental design. **(B)** Quantification of IFNγ concentration in the serum of IFNγ (n=8) and CTL (n=7) mice performed by ELISA assay, 14 days post-injury. Mann-Whitney U test, means ± SEM. **(C)** Quantification of *CIIta* (*left*) and *Mhc-II* (*right*) gene expression by RT-qPCR performed on Tibialis Anterior (TA) injured muscles from IFNγ (n=8) and CTL mice (n=8), 14 days post-injury. Mann-Whitney U test, means ± SEM. **(D)** Representative immunofluorescence images of LAMIMIN (white), MHC-II (red), and nuclei (DAPI, blue) performed on IFNγ and CTL injured TAs, 14 days post-injury. **(E)** Representative immunofluorescence images of LAMININ (green), CD68 (red), and nuclei (DAPI, blue) performed on IFNγ and CTL injured TAs, 14 days post-injury. **(F)** Quantification of the percentage of CD68+ macrophages per area performed on IFNγ (n=7) and CTL (n=8) injured TAs, 14 days post-injury. Mann-Whitney U test, mean± SEM. **(G)** Quantification of *Inos* (*left*) and *Tgfβ* (*right*) gene expression performed by RT-qPCR performed on IFNγ (n=7) and CTL (n=8) injured TAs, 14 days post-injury. Mann-Whitney U test, means ± SEM. **(H)** Representative images of collagen deposit by Sirius Red staining (*left*) and quantification of the percentage of connective tissue area (*right*) performed on injured TAs from IFNγ and CTL mice, 14 days post-injury. Mann-Whitney U test, means ± SEM. **(I)** Representative images of Hematoxilin eosin staining (*left*) and quantification of the percentage of adipocytes invasion by bodipy staining (*right*) performed on injured TAs from IFNγ (n=7) and CTL (n=6) mice, 14 days post-injury. Mann-Whitney U test, means ± SEM.

Altogether, our data showed that continuous high levels of circulating IFNγ triggers the IFNγ- signature in injured muscles, with increased macrophage infiltrate, endomysial fibrosis, and adipose involution, mimicking most of the molecular and cellular features previously observed in IBM muscle *(Figure 1)*.

### Systemic elevation of IFNγ delays myofiber regeneration in mice

Next, we sought to identify the mechanisms of action underlying the deleterious effect of IFNγ in IBM. IFNγ is a potent pro-inflammatory cytokine secreted by M1 macrophages, which are known to delay the kinetics of muscle differentiation^26^. Then, we examined whether the upregulation of the IFN-II signature affects myogenesis *in vivo (Figures 3A)*. The number of regenerating embryonic MyHC-positive (eMHC+) myofibers in IFNγ and CTL mice was monitored. eMHC protein is typically expressed from 2-3 days post-injury, then completely cease to be expressed between 7 and 14 days^27^. Although we did not observe a difference at day 7, there was a significant increase of the density of eMHC+ fibers in TAs of mice treated with IFNγ at day 14 (*Figure 3B, C*). This phenotype was associated with a reduction of regenerating TA muscle weight *(Figure 3D)* and myofiber size *(Figure 3E)* in IFNγ mice. In contrast, non-injured *Gastrocnemius* muscle did not show any change in myofiber size under systemic IFNγ delivery *(data not shown)*, indicating that the observed atrophic phenotype of TA muscle was linked to the phenomenon of post-lesional repair. While multi-nucleated fibers were present abundantly in CTL mice, the number of centralized nuclei per fiber was significantly decreased at 14 days post-injury in IFNγ-exposed regenerating TAs *(Figure 3F)*, indicating an overall alteration of myoblastic cell fusion to growing regenerating myofibers under systemic IFNγ exposure. We next examined whether sustained IFNγ release affect myogenic properties of muscle satellite cells (MuSCs), which are known to trigger muscle repair. Indeed, IFNγ promotes the classical activation of macrophages, which delay myogenesis by extending the proliferative period of myoblasts before they enter myogenic differentiation^26,28^. However, seven days post-injury, the density of PAX7+ cells *(Figure 3G, H)* as well as the proportion of proliferating PAX7+KI67+ cells *(Figure 3H)* was decreased in regenerating TAs of IFNγ mice, not suggesting a central role for IFNγ- induced M1 activation in post-injury muscle atrophy. Thus, systemic IFNγ stress triggers sporadic and muscle-specific loss of MuSC density and function, participating muscle repair delay.

**Figure 3.**
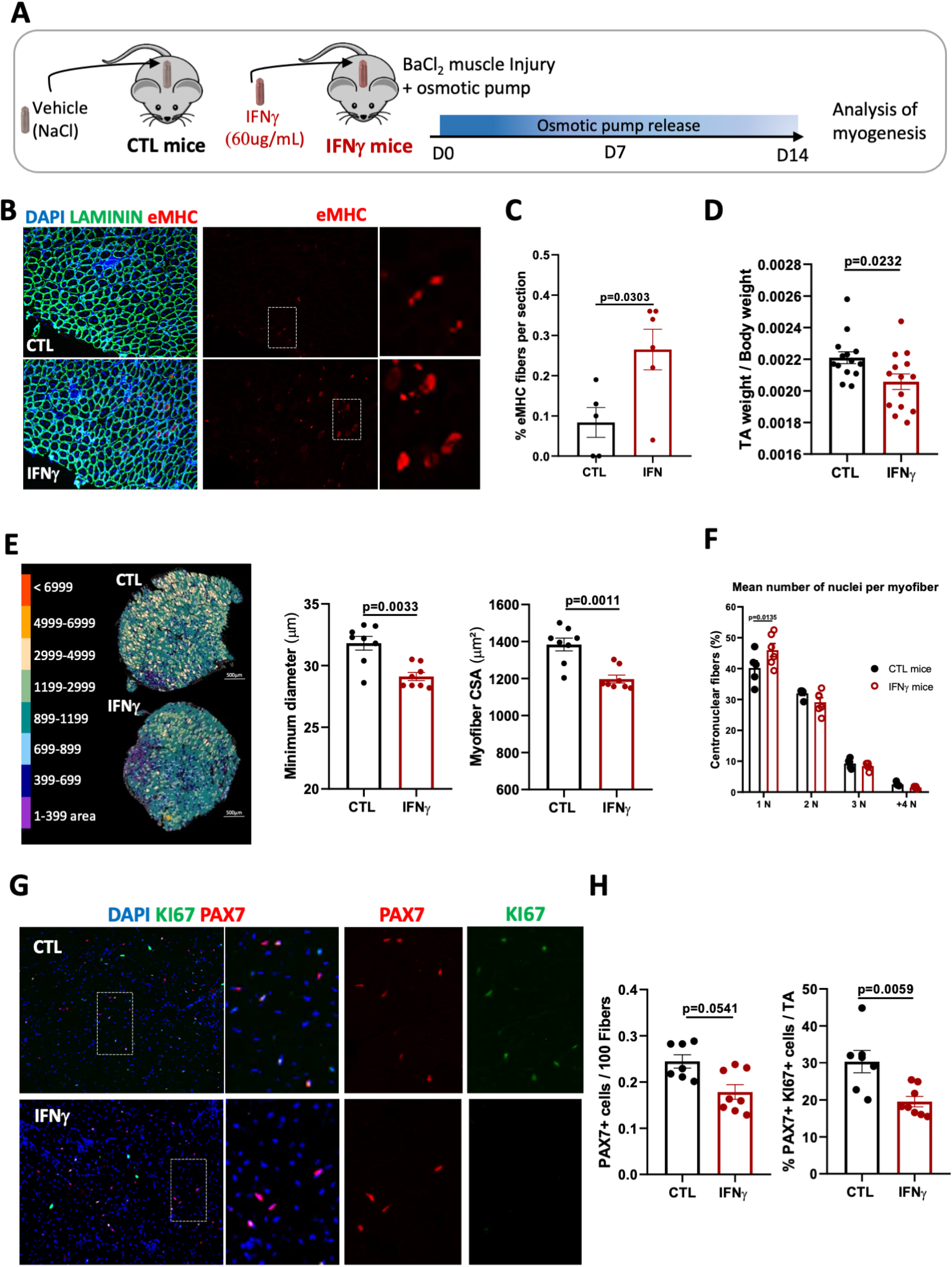
Systemic elevation of IFNγ delays myofiber regeneration in mice. **(A)** Experimental design. **(B)** Representative immunofluorescence images of LAMIMIN (green), embryonic isoform of myosin heavy chain, eMHC (red), and nuclei (DAPI, blue) performed on IFNγ and CTL injured TAs, 14 days post-injury. **(C)** Quantification of the percentage of eMHC+ fibers per area performed on IFNγ (n=8) and CTL (n=8) injured TAs, 14 days post-injury. Mann-Whitney U test, mean± SEM. **(D)** Quantification of TA muscle mass quantification per body weight at 14 days post-injury in IFNγ (n=14) and CTL (n=14) mice, 14 days post-injury. Mann-Whitney U test, means ± SEM. **(E)** Myofiber size analyses based on laminin immunostaining and automated analysis, as previously described^55^ (left), minor diameter, and cross-section-area (CSA) quantifications performed on IFNγ (n=8) and CTL (n=8) injured TAs. Mann-Whitney U test, means ± SEM. **(F)** Quantification of nuclei number by centronuclear fibers per area performed on IFNγ (n=7) and CTL (n=6) injured TAs, 14 days post-injury. Two-Way ANOVA, Sidak’s multiple comparison. **(G)** Representative immunofluorescence images of PAX7 (red), proliferating marker KI67 (green), and nuclei (DAPI, blue) performed on IFNγ and CTL injured TAs, 7 days post-injury. **(H)** Quantification of PAX7+ cells per 100 fibers (left) and percentage of proliferating MuSC PAX7+ Ki67+ per PAX7 total (right) performed on IFNγ (n=8) and CTL (n=7) injured TAs, 7 days post-injury. Means ± SEM, Mann-Whitney U test.

### IFNγ directly repress the proliferation, activation and fusion of myogenic cells

To directly address the impact of increased IFNγ levels on the myogenic capacities of MuSCs, we exposed human muscle progenitor cells (MPCs) to IFNγ (2×10^3^ U/mL) *(Figure 4A)*. MPCs were isolated and purified from CTL human deltoid muscles, as previously described^29^. After 72h, IFNγ-exposed MPCs expanded less in culture than CTL MPCs *(Figure 4B)*. This reduced proliferation was confirmed using the Ki67 proliferation marker that showed a decrease of Ki67 expression in IFNy-treated MPCs at the mRNA (*Figure S3A*) and protein level *(Figure 4C, D*) up to 10 days of exposure. Of note, the decrease of cell proliferation, also illustrated by a decrease in ATP content *(Figure S3B*), was not due to cell death since IFNy binding did not increase the plasma membrane permeability *(Figure S3C)*. Besides proliferation, IFNγ also inhibited the migration capacity of MPCs *in vitro*, as evaluated by the scratch test after 48h of exposure *(Figure S3 D, E)*. Moreover, the capacity of MPCs to activate *in vitro* was inhibited by IFNγ, as assessed by the reduced number of PAX7^**+**^ MYOD^**+**^ cells *(Figure 4E)*. To further characterize the direct impact of IFNγ exposure on MuSC fate, we performed complementary experiments using *ex vivo* floating murine myofibers that offer the advantage to retain MuSCs within their niche^30^. Using this method, we confirmed that *in vitro* IFNγ exposure decreased the percentage of activated Pax7^**+**^ MYOD^**+**^ cells towards an increase of Pax7^**+**^ MYOD^**-**^ self-renewing cells after 48h *(Figure 4F)*. In human MPCs, the decreased activation upon IFNγ stimulation was accompanied by a decrease in the mRNA level of myogenic differentiation markers, *Myogenin (MyoG)* and *Myf6 (Figure S3F)*, demonstrating an overall impairment of both the activation and differentiation potential of myogenic progenitors by IFNγ.

**Figure 4.**
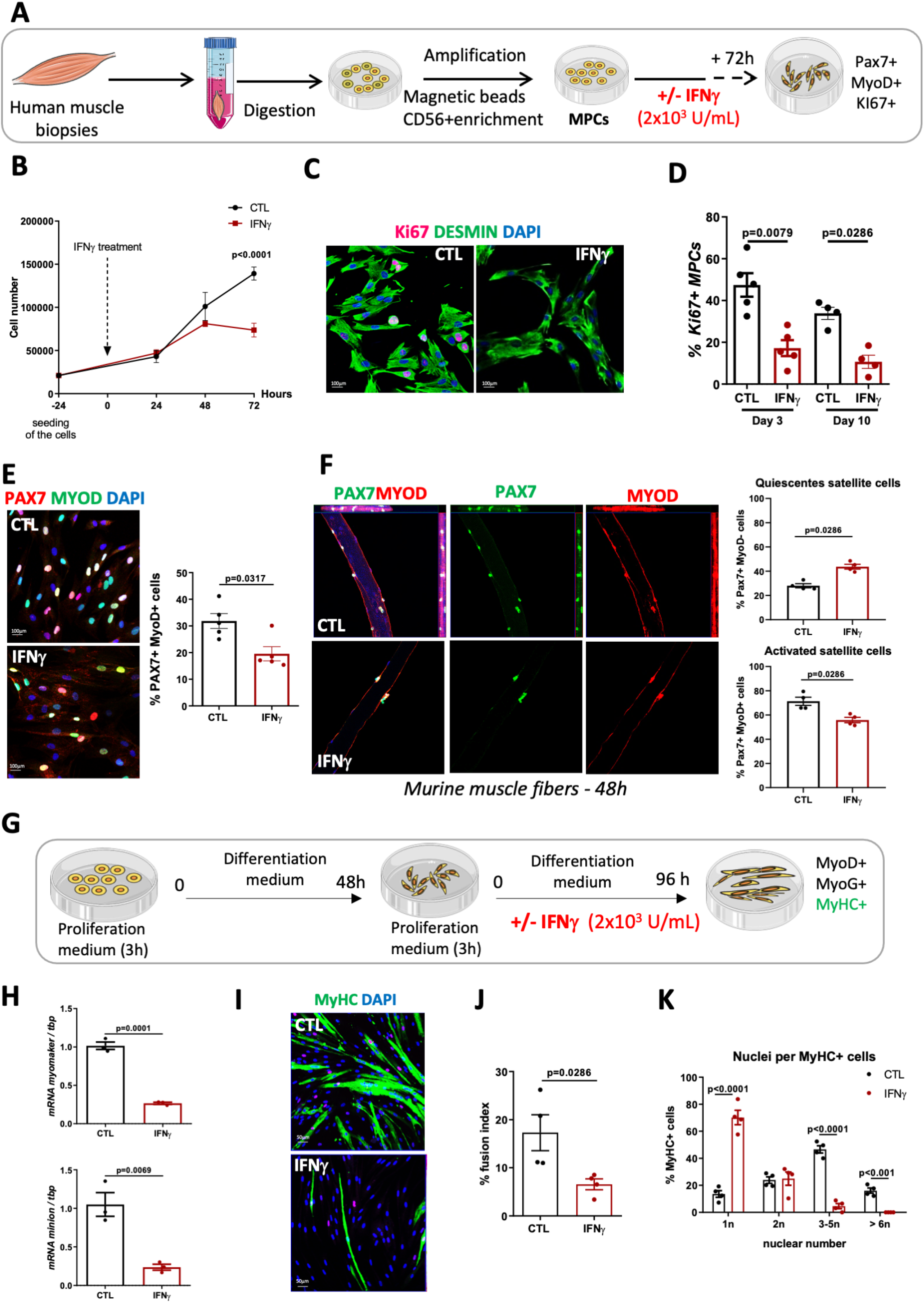
IFNγ directly repress the proliferation, activation and fusion of myogenic cells. **(A)** Experimental design for *in vitro* studies (n=4). **(B)** Human myogenic progenitor cells (MPC) growth curve, exposed or not to IFNγ for 72h. Two way-ANOVA, Sidak’s multiple comparisons test. **(C)** Immunofluorescence staining of DESMIN (green), Ki67 (pink), and nuclei (DAPI, blue) in CTL and IFNγ exposed-MPC. **(D)** Quantification of proliferating DESMIN+ Ki67+ MPCs percentage, at 3 and 10 days. Means ± SEM Mann-Whitney U t test. **(E)** Immunofluorescence staining of MYOD (green), PAX7 (red), and nuclei (DAPI, blue) (*left*) and PAX7+ MYOD+ cell (*right*) percentage performed on CTL and IFNγ exposed-MPC, at 72 hours. Mann-Whitney U test, means ± SEM. **(F)** Immunofluorescence staining of PAX7 (green), MYOD (red), and nuclei (DAPI, blue) (*left*) and quantification of the of PAX7+ MYOD+ cells and PAX7+ MYOD-(*right*) percentages performed on floating murine myofibers, exposed or not to IFNγ for 48hours. Mann-Whitney U test, means ± SEM. **(G)** Experimental design for the differentiation and fusion studies. **(H)** Quantification of *myomaker* (up) and *minion* (bottom) fusogene expression by RT-qPCR performed on CTL and IFNγ exposed-MPCs, at 96 hours. Mann-Whitney U test, means ± SEM. **(I)** Immunofluorescence staining of MyHC (green) and nuclei (DAPI, blue) performed on CTL and IFNγ exposed-myocytes and myotubes, for 96 hours. **(J)** Fusion index of synchronized CTL and IFNγ exposed-MPCs after 96h of differentiation. Mann-Whitney U test, means ± SEM. **(K)** Quantification of myotubes size, number of nuclei per MyHC positive cells. Mann-Whitney U test, means ± SEM.

To decipher whether sustained IFNγ delivery induces muscle atrophy *in vivo* by impairing muscle progenitor differentiation solely or by impacting also the fusogenic capacity of myogenic progenitors, we exposed human MPCs in differentiation medium for 96 hours at low density to synchronize them in a differentiated state (mainly, MYOD^**+**^ and MYOG^**+**^ cells) *(Figure 4G)*. Then, differentiated MYOD^**+**^MYOG^**+**^ cells were split and seeded at high density to allow them to fuse. Using this strategy, we showed that IFNγ exposure decreased in differentiated MPCs the transcript level of *Myomaker* and *Myomerger* (*Figure 4H*), the two main fusogenic proteins. As a consequence, the fusion index of MPCs was strongly decreased upon IFNγ treatment *(Figure 4I, J)* and IFNγ–exposed MPCs formed fewer and smaller myotubes *(Figure 4K)* compared to CTL MPCs. Overall, these results witnessed that chronic IFNγ exposure slows down the differentiation as well as the fusion capacity of myogenic progenitors, participating in the atrophic muscle fiber phenotype observed in injured and IBM muscles *in vivo*.

### Elevated IFNγ level increases premature cellular senescence

Replicative senescence is associated with impaired differentiation capacity of human myogenic progenitor cells^31,32^. Since IFNγ induces cell senescence in many cell types^33,34^, we examined whether chronic IFNγ exposure may trigger premature senescence of myogenic progenitors *in vitro* and *in vivo*. Treatment of human MPCs with IFNγ led to senescent-associated morphological changes, including a typical flat and enlarged cellular shape as compared to CTL MPCs *(Figure 5A,B)*. Using busulfan as a positive control for senescence induction^35^, we observed similar levels of SAβGal-positive cells in IFNγ- and busulfan-treated MPCs (*Figure 5D,E*). Interestingly, IFNγ-exposed MPCs showed an increase of *p16* mRNA level *(Figure 5C)* and SAβGal staining (*Figure 5D,E*), two senescence-associated markers^36^. To verify whether IFNγ signature is also associated with cell senescence in pathology, we investigate whether MHC-II+ MPCs isolated from IBM/ASS patients with high IFN-II signature showed evidence of premature senescence compared to CTL MPCs *(Figure 5G-K)*. We found that the MPCs from MHC-II+ muscles proliferated less rapidly than CTL MPCs, as evaluated by the decrease in the cell doubling time all along the passages *(Figure 5F, G). In vivo*, the ratio of PAX7^**+**^Ki67^**+**^ myogenic progenitors was lower in IBM than in other DIMs muscles (*Figure 5H, I)*, suggesting that the proliferative capacity of MPCs in IBM muscles is reduced. The decreased proliferation of muscle stem cells in IBM, associated with an upregulation of *p16* transcripts *(Figure 5J)* and senescence-associated pathways *(Figure 5K)*.

**Figure 5.**
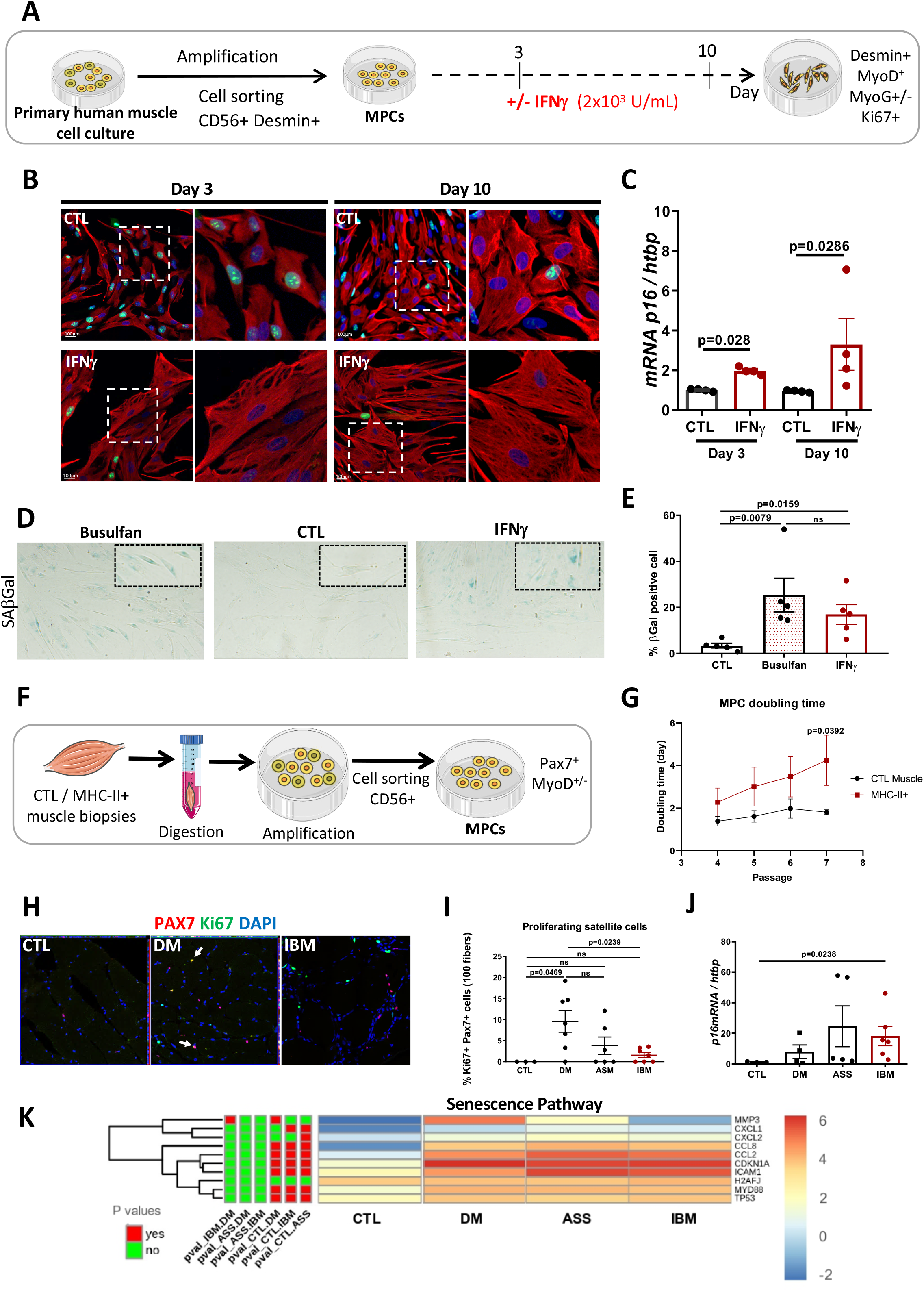
Elevated IFNγ level increases premature cellular senescence. **(A)** Experimental design (n=4 independent experiments per condition) **(B)** Representative immunofluorescence images of DESMIN (red), KI67 (green) and nuclei (DAPI, blue) performed on MPCs exposed or not to IFNγ for 72 hours and 10 days. **(C)** Quantification of *p16* gene expression by RT-qPCR performed on CTL and IFNγ exposed-MPCs, at 72 hours and 10 days. Mann-Whitney U test, means ± SEM. **(D)** Representative images of SAβGal staining performed on MPCs, exposed or not to IFNγ for 10 days. **(E)** Quantification of SAβGal positive cells at 10 days in CTL, IFNγ and Busulfan (positive control of senescence) conditions. Means ± SEM, Unpaired T-test. **(F)** Experimental design obtain purified CD56+ MPCs from healthy donor and MHC-II+ muscle from IBM/ASS patients. **(G)** Quantification of the doubling time (day) for several passages of MPCs from healthy donor (CTL) and muscle with strong myofiber MHC-II expression (MHC-II+). Two-way ANOVA, Sidak’s multiple comparison **(H)** Representative immunofluorescence images of PAX7 (red), proliferating marker KI67 (green), and nuclei (DAPI, blue) performed on CTL, DM, ASS and IBM muscle sections. **(I)** Quantification of the percentage of proliferating MuSCs PAX7+ KI67+ cells. Means ± SEM. Mann-Whitney U test. **(J)** Quantification of *p16* gene expression by RT-qPCR performed on CTL (n=3), DM (n=4), ASS (n=5) and IBM (n=6) muscle. Means ± SEM, Unpaired T-test. **(K)** Heatmap senescence pathway with the normalized reads per gene of CTL (n=5), DM (n=5), ASS (n=5) and IBM (n=4) RNA-seq data.

Together, these data confirm muscle stem cell dysfunction in IBM patients associated with premature cellular senescence, impaired regenerative capacity, and abnormal muscle overexpression of IFNγ.

### The deleterious effects of IFNγ on myogenic cells are mediated by JAK-STAT activation

IFNγ triggers immune responses through activation of JAK1/2 receptor and induction of STAT1 phosphorylation that activates MHC-II expression on the cell surface via the CIITA transactivator^10,11^. We thus evaluated whether IFNγ controls MHC-II expression in MPCs through JAK-STAT signaling. By western-blot analysis, we showed that IFNγ increased in human MPCs the activation of JAK-STAT pathway, as assessed by the increased expression of both STAT1 protein and its phosphorylated form after 6 days of IFNγ exposure *(Figure 6A, B)*. This result correlated the transcriptomic studies performed in IBM muscles, showing an upregulation of STAT1 and its downstream target genes (CIITA, HLA-DR, -DO) *(Figure S1C)*. Activation of the JAK-STAT pathway under IFNγ was confirmed by the parallel increase of *CIIta* transactivator and *Mhc-II* gene expressions in human IFNγ-treated MPCs *(Figure 6C)*. As expected, overexpression of MHC-II protein under IFNγ was located mainly at the surface of human MPCs, whatever their myogenic state, *i*.*e*., myoblast, myocytes and myotubes *(data not shown)*.

**Figure 6.**
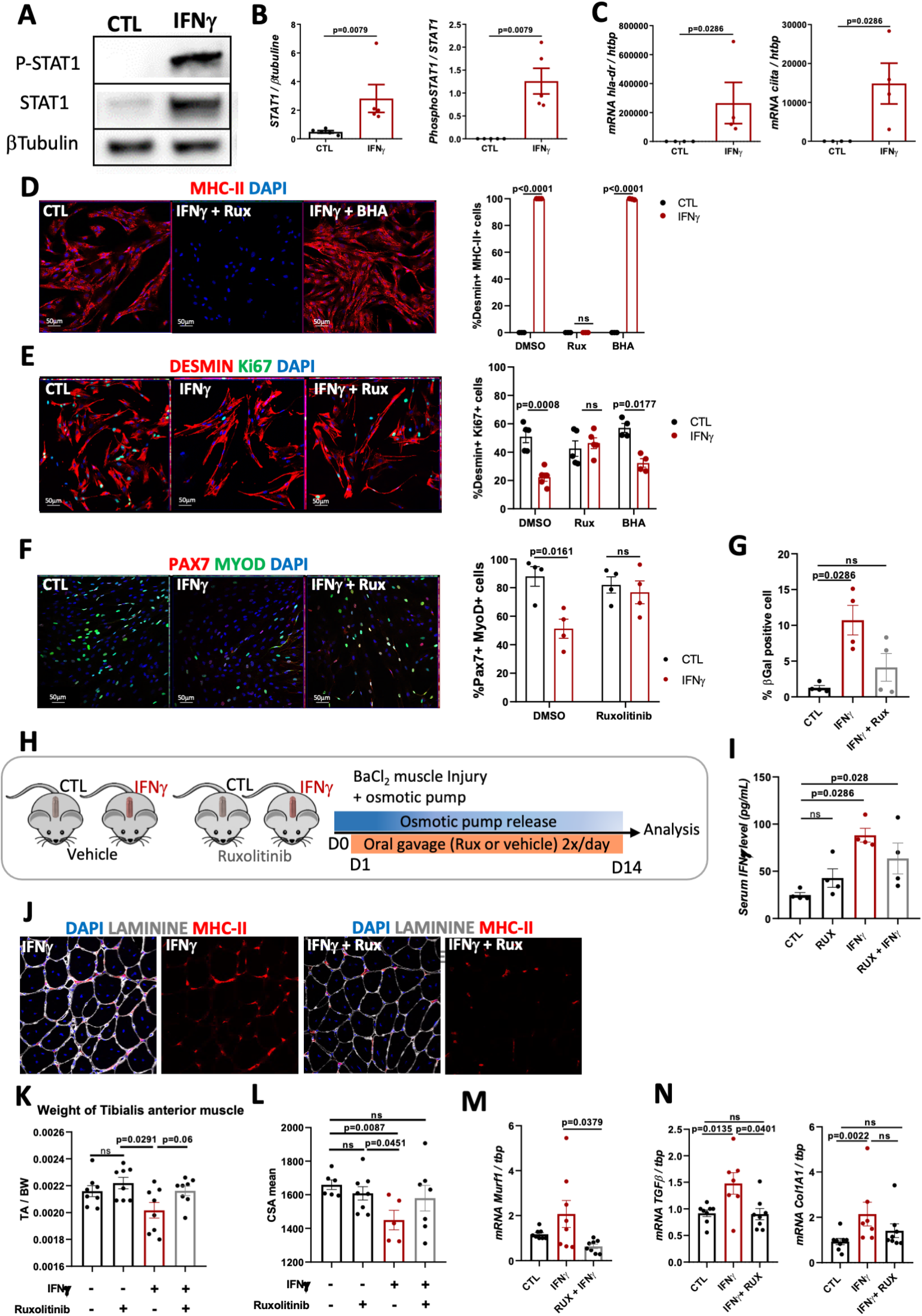
**(A)** Immuno blot for Phospho-STAT1, STAT1 and βTubulin (left); human MPCs after 6 days of culture ± IFNγ exposure (n=5). Uncropped blots in Source Data. **(B)** STAT1 and Phospho-STAT1 signal intensity. Means ± SEM, Mann-Whitney U test. **(C)** C*IIta* (left) *and Hla-Dr* (right) gene expressions in human CTL and IFNγ-exposed MPCs for 6 days (n=4). Means ± SEM Mann-Whitney U test. **(D)** Immunofluorescence staining (left) of MHC-II (red), and nuclei (DAPI, blue) in CTL, IFNγ-exposed MPC and with or without specific inhibitors, Ruxolitinib anti JAK-STAT1, BHA anti-oxydative stress. MHC-II+ cells (right) quantification at 72 hours. One-way ANOVA, Dunn’s multiple comparison. **(E)** Immunofluorescence staining (left) of DESMIN (red), KI67 (green) and nuclei (DAPI, blue) in CTL, IFNγ-exposed MPC and IFNγ-exposed MPC with ruxolitinib treatment at 72h. KI67+ DESMIN+ MPCs quantification at 72 hours (right). Means ± SEM Mann-Whitney U test. **(F)** Immunofluorescence staining (left) of PAX7 (red), MYOD (green) and nuclei (DAPI, blue) in CTL MPCs, IFNγ-exposed MPCs and IFNγ-exposed MPCs with ruxolitinib treatment at 72h. Pax7+ MyoD+ quantification at 72 hours. Means ± SEM Mann-Whitney U test. **(G)** SAβGal positive cells quantification at 5 days in CTL MPCs, IFNγ-exposed MPCs and IFNγ-exposed MPCs with ruxolitinib. Means ± SEM Mann-Whitney U test. **(H)** Experimental design of *in vivo* studies, 4 groups of transplanted wild-type mice with subcutaneous osmotic pump releasing IFNγ or NaCl for 14 days. Groups received ruxolitinib (Rux) or solvent by oral gavage twice daily for 14 days. **(I)** Serum IFNγ concentration in IFNγ (n=8), CTL (n=7), Rux and Rux+IFNγ mice performed by ELISA assay, 14 days post-injury. Mann-Whitney U test, means ± SEM. **(J)** Immunofluorescence staining of LAMININ (white), MHC-II (red), and nuclei (DAPI, blue) in CTL and IFNγ injured TA muscle. **(K)** TA muscle mass at 14 days post-injury in IFNγ, CTL, Rux and Rux+IFNγ mice (n=8 per group), 14 days post-injury. Mann-Whitney U test, means ± SEM. **(L)** Cross-section-area (CSA) quantifications in IFNγ (n=8), CTL (n=8), Rux and Rux+IFNγ injured TAs. Mann-Whitney U test, means ± SEM. **(M)** *Murf1* atrogene expression in CTL, IFNγ, Rux and Rux+IFNγ injured TAs, 14 days post-injury. Means ± SEM Mann-Whitney U test. **(N)** *TGFβ* (left) and *Col1A1* (right) expressions in CTL, IFNγ, Rux and Rux+IFNγ injured TAs, 14 days post-injury. Means ± SEM Mann-Whitney U test.

We then analyzed whether the activation of the JAK-STAT pathway is responsible for the deleterious effects of IFNγ on myogenic cell functions. For this purpose, human MPCs were treated with IFNγ in the presence or absence of a specific JAK1/2 inhibitor, ruxolitinib (*Rux*), in comparison with butylated hydroxyanisole (BHA) a molecule which being able to block reactive oxygen species (ROS)-mediated senescence. Ruxolitinib treatment completely abolished IFNγ-mediated MHC-II expression *(Figure 6D)* and rescued the inhibitory effects of IFNγ on the proliferation *(Figure 6E)*, the activation *(Figure 6F)* and the premature senescence *(Figure 6G)* of human MPCs, as assessed by the ratio of proliferating DESMIN^**+**^KI67^**+**^, activated PAX7^**+**^MYOD^**+**^ and βGAL^**+**^ cells, respectively. BHA did not prevent the repressive effect of IFNγ on myoblasts proliferation, indicating that this effect did not depend on ROS production. Our data therefore demonstrate that IFNγ alters the myogenic function of human MPCs *in vitro*, via the activation of JAK-STAT signaling.

### JAK-STAT inhibitor reverses the deleterious effects of IFNγ on muscle regeneration

To determine whether JAK1/2 inhibitor can prevent IFNγ-mediated muscle lesions in injured muscles *in vivo*, we treated injured mice with either ruxolitinib (9.6 mg/kg/day) or solvent (controls, CTL), by oral gavage twice daily for 14 days *(Figure 6H)*. Ruxolitinib treatment did not affect the serum concentrations of IFNγ *(Figure 6I)*. In contrast, ruxolitinib significantly reduced the expression of muscle MHC-II, confirming the efficacy of the treatment to antagonize JAK1/2 signaling and the IFNγ signaling activation *in vivo (Figure 6J)*. Importantly, we showed that ruxolitinib did not affect muscle repair in CTL mice but specifically restored muscle regeneration capacity in IFNγ-exposed mice (*Figure 6J-N*). Indeed, ruxolitinib-treated IFNγ mice showed similar TA muscle weight and myofiber size compared to CTL mice *(Figure 6K-L)*. In line with these results, the expression of *Murf1* atrogene was increased in IFNγ mice compared to CTL and decreased with ruxolitinib treatment *(Figure 6M)*. Furthermore, ruxolitinib significantly decreased *Col1A1* and *TGFβ* gene expressions in IFNγ mice which are markers for developing fibrosis *(Figure 6N)*. Our results, therefore, demonstrated that high IFNγ level exerts deleterious effects on regenerating muscles *in vivo* involving muscle atrophy and fibrosis via the JAK-STAT signaling pathway. These properties were antagonized *in vivo* by the ruxolitinib.

## Discussion

IBM is an idiopathic and slowly progressing disease affecting muscle function, which does not benefit from well-admitted animal models to investigate its pathogenesis^1,37^. Typical histological features associate myonecrosis, protein aggregates within myofibers, and endomysial inflammation. We confirmed that DIM strikingly differ according to their IFN signatures. IFN-I signature is the highest in DM, and IFN-II in ASS and IBM, IBM and ASS displaying similar levels of IFN-I/II-stimulated genes. Our results highlight the importance of gene selection to qualify IFN signatures which may explain discrepancies that can be noted between studies^7,38^. The MHC-II expression was also characterized at the protein level by immunohistochemistry and clearing method that confirmed the localization of MHC-II protein at the sarcolemma and therefore the myofiber response to IFNγ.

In IBM, the combination of a very long and slow course of the disease with the presence of IFNγ- secreting CD8 T-cells in endomysium, necessarily leads to a strong and sustained exposure of myofibers to IFNγ, without commensurate with what is observed in other DIMs, and makes plausible the direct implication of IFNγ in myofiber alterations. In IFNγ mice, myofibers abnormally express MHC-II expression. In case of muscle injury, increased circulating IFNγ was associated with delayed repair, endomysial fibrosis and adipocyte accumulation. These muscle damages are in line with the typical histological characteristics of IBM muscles. These data are reminiscent of the cardiomyopathy developed in SAP-IFN-γ5 mice, with atrophy and fibrosis^39^. A typical hallmark of IBM muscle is the presence of amyloid depositions that we did not observe in IFNγ mice. Interestingly, the culture of myoblasts in the presence of IFNγ and IL1β for 48 hours is accompanied by the formation of amyloid aggregates^40^. Our RNAseq analysis showed an upregulation of others proinflammatory cytokines in IBM muscles. It is possible that a synergic effect of proinflammatory cytokines in muscle could be promote muscle IBM-like features, such as amyloid deposits.

The uninjured muscles showed neither MHC-II positive myofibers, atrophy nor fibrosis *(data not shown*), suggesting that the development of IBM requires the combination of repeated muscle injuries with deregulated immune response, leading to protracted release of IFNγ in the vicinity of myofibers, and secondary to the occurrence of degenerative changes. These data are in line with results obtained using SAP-IFN-γ5, since authors did not observe a pathogenic phenotype in uninjured quadriceps muscle^39^. In case of muscle injury, systemic elevation of IFNγ was also associated with an increase of macrophage population. This increase likely corresponds to the post-myonecrosis recruitment of macrophages and reflects the delay in muscle regeneration of IFNγ mice. This cytokine is able to stimulate M1 macrophages and favor the proliferation of myoblasts^41^. However, our data showed that IFNγ directly impairs satellite cell proliferation and differentiation, and promotes myofiber atrophy *in vivo*. The systemic elevation of IFNγ increased fibrosis and adipocyte accumulation in mice muscle, both features being characteristic of the fibroadipogenic involution process observed in chronically diseased muscles. Finally, our results showed that, as much a transient increase of IFNγ and activation of M1 macrophages is necessary for myofiber regeneration, as much the protracted increase of IFNγ in myofiber microenvironment is profoundly deleterious for muscle tissue repair.

IFNγ signaling leads to the induction of CIITA transactivator, which combines with promotors to launch MHC-II expression^42^. In the murine myogenic cell line C2C12, CIITA represses myogenesis by inhibiting myogenin^43,44^. We also showed that IFNγ stimulation induces the myogenic cells senescence in addition to myogenesis repression. In accordance with these experimental findings, we showed that MPCs from muscles with strong myofiber expression of MHC-II proliferated less rapidly than those from non-diseased muscles, regardless of IFNγ presence in the culture medium, indicating a long-term effect of IFNγ on MPC growth capacity. Likewise, the proportion of proliferating satellite cells in muscle biopsy samples from IBM patients was lower than in DM patients. Transcriptomic analysis confirmed the senescence pathway activation in IBM muscle. IBM is also characterized by a defect in the functionality of satellite cells. Thus, our data suggest that IFNγ could mediate senescence activation through p16 in IBM muscle. Multiple stimuli are able to induce senescence, which is regulated mainly by the tumor suppressors p16, p53, Rb, as well as the cyclin-dependent kinase inhibitors^45^. Inflammatory stimuli such as TNFα binding has been shown to up-regulate IFNγ signature and mediate STAT1-mediated senescence in HUVEC cells^46^.

Transcriptomic studies also showed activation of the STAT1 pathway, with increased STAT1 and its phosphorylated form (P-STAT1) in IBM patients. Therefore, we aimed at determining the relevance of using JAK/STAT inhibitors to repress the deleterious effect of chronic IFNγ up-regulation in regenerating muscles. Ruxolitinib is an oral inhibitor of the JAK-STAT pathway and is already tested in clinics to prevent IFN type I dependent cytotoxicity^47–49^. In our hands, the pharmacological inhibition of JAK1/2 completely blocked the IFNγ signaling pathway in MPCs, reflected by MHC-II and CIITA expression and restored their normal proliferation and activation upon IFNγ exposure *in vitro*. Interestingly, it was shown that the reduction of muscle regenerative capacities and satellite cell senescence in aged mice was associated with the upregulation of JAK-STAT signaling targets. In addition, the pharmacological inhibition of Jak2 or Stat3 stimulates satellite stem cell divisions and enhances the repopulation of the satellite cell niche^50^ supporting the potential therapeutical interest of JAK inhibition in IBM.

*In vivo*, we validated the effects of Ruxolitinib on muscle phenotype developed in IFNγ-treated mice. Ruxolitinib dosage (9.6 mg/kg/day), was largely below what is used to cancer treatment in mice (50mg/kg/day) cancer^51^, suggesting that our findings might be of use for preclinical studies. IFNγ- induced muscle atrophy was rescued by Ruxolitinib treatment, which also dampened the expression of fibrosis markers Col1A1, and TGFβ. Thus, targeting JAK1/2 can prevent major deleterious effects of IFNγ on muscle without generating any obvious adverse effects^49^.

Recently, a clinical assay showed some beneficial effects of rapamycin, mTOR pathway inhibitor in IBM patients^52^. Our RNA sequencing analysis indicated a downregulation of mTOR signaling pathway *(Figure S1D)* and specifically the number of mTOR transcript is unchanged between CTL and IBM muscles *(Figure S1E)*, suggesting that mTOR pathway regulation is not directly involved in IBM. Moreover, it must be noted that muscle-specific mTOR knock-out mice present severe myopathy^53^ and muscular adverse effects of long-term rapamycin treatment have been reported in transplanted patients^54^.

In conclusion, our data extended the characterization of IBM pathogenesis with a decrease in the regenerative capacities of muscle satellite cells, and that the up-regulation of IFNγ signaling is one of hallmark in IBM pathogenesis. Ectopic IFNγ overexpression recapitulates typical histological feature of IBM muscles such as myofiber atrophy, fibrosis, adipocytes invasion and cell senescence by activating the JAK/STAT pathway. Moreover, we provided experimental evidence supporting the efficiency of the JAK/STAT inhibitor ruxolitinib to counteract the deleterious effect of IFNγ in interferonopathies affecting muscle phenotype such as IBM. JAK-STAT could be a new therapeutic target, suggesting that ruxolitinib or others JAK-STAT inhibitors may be of use for IBM patients.

## Materials and methods

### Patients and muscle samples

Muscle samples were collected by FJA, SS and EM at Mondor hospital from patients undergoing muscle biopsy for diagnostic purposes (*Table 1*). Samples were obtained from deltoid muscles. All human specimens were collected with informed consent and procedures approved by IRB (Henri Mondor Biological Resource Platform: registration number DC-2009-930, French Ministry of Research). Muscle biopsy samples were conventionally processed for myopathology^6^ with immunoperoxidase staining of MHC-I and MHC-II antigens, complement membrane attack complex (C5b-9), CD56/NCAM (myofiber regeneration), CD68, CD3, CD4, CD8, and CD20 (leukocyte subsets) performed on 7 µm-cryosections (references in^6^). All biopsy specimens were reviewed blindly for clinical and MSA data by the FJA, SS and EM. ENMC criteria were used to diagnose dermatomyositis (DM)^56^, criteria published in 2014 by Lloyd et al. to diagnose inclusion body myositis (IBM)^57^, and Troyanov classification to diagnose overlap myositis^58^ the ASS subset of which being defined by detection of circulating anti-synthetase auto-antibodies. Controls (CTL) were patients presenting with chronic myalgias, but no definite neuromuscular pathology after diagnostic work-up including muscle biopsy (histologically normal muscle).

### Isolation of muscle progenitor cells from human deltoid muscle biopsy

Human muscles were dissociated and digested with pronase enzyme *(1*.*5mg/mL; protease from streptomyces griseus P5147-5G Sigma)* in *DMEM* at 37°C for 20 min and passed through a 100 mm cell strainer (repeated 4 times). Then, stopped the digestion activity by FBS (30%). Centrifugate cells surnageant 20 min at 1600 rpm. Cells were seeded onto 5 flasks coated with gelatin 5cm2 in F12 medium (*Life Technologies, Gibco® 31765-027*) with 20% FBS (*Dutscher; CAT S1810-500* ; *lot S11307S1810*), 0.2% Vitamines, 1% AANE, 1% P/S and ultroserG 1% (*Life sciences 15950-017*).

### Muscle primary cell culture

After cellular amplification, the cells are sorted with CD56 microbeads *(Miltenyi, 130-050-401)* two times, the efficiency is 90-95% CD56+ cells. Human muscle progenitor cells (MPC) were cultured in F12 medium with 20% FBS, 0.2% Vitamines, 1% AANE, 1% P/S in a humidified atmosphere at 37°C and 5% CO2.

### IFNγ and Inhibitor treatments

Cells were treated with human IFNγ (*Human IFN-Y1b, 100ug, ref 130-096-484 Miltenyi*). Cells were stimulated with several IFNγ concentration, and the majority of experiments was made with 2×10^3^ U/mL (25ng/mL) of IFNγ. Cells were harvested for RNA or protein at defined time points after the IFNγ exposure. For acute exposure, IFNγ was added just one time 24h after the seeding and the medium was not replace for 72 hours. Moreover, for sustained exposure, IFNγ was added every day in 50% of medium for 72h or 10 days. Some experiments including Ruxolitinib (Rux), selective inhibitor of JAK1/2 (10uM) *(Invivogen)*, and Butylated hydroxylanisole (BHA), anti-reactive oxygen species (100uM) exposure for 72h. At least four independent experiments were assayed for each data point.

### Muscle progenitor cell proliferation

For proliferation experiments, primary human MPC were seeded at 6000 cells/cm2 in culture medium. Next day, the medium was changed by F12 with 10% FBS, 1% P/S supplemented with or without IFNγ treatment during 3 or 10 days with split when muscle cells obtain 80% confluence.

### Myogenic differentiation

For differentiation experiments, primary human MPC were seeded at confluence density (20 000 cells/cm^2^) in culture medium. After 2 days, when the cells confluence reached 80%, the culture medium was changed for differentiation medium (F12, 2% Horse serum, 1% P/S) supplemented with or without IFNγ treatment during 3 days. Fusion index is expressed as the ratio of the nuclei number in myocytes with two or more nuclei versus the total number of nuclei.

### Senescence analysis

For the positive control, cellular senescence was induced by Busulfan drug (*B2635-10G Sigma*). The cells were exposed to varying concentrations of Busulfan (50uM and 150 uM) for 24 h. The cells were washed twice with PBS 1X to remove drug and reseeded in fresh medium for 5 or 10 days.

### SA-β-gal Staining

Cells were fixed in a solution of PBS, 1% PFA, 0.2% glutaraldehyde for 5min at RT. Samples were washed in PBS pH7 2X 10min and incubated for 30 min in PBS pH6 and further incubated in an X-gal solution (4mM K3Fe(CN)6, 4mM K4Fe(CN)6, 2mM MgCl2, 0.02% NP-40 (Igepal) and 400 mgml1 X-gal (15520-018 Sigma) in PBS pH6, at 37 °C ON for cells and 2×24 h for sections, according to the publication from^59^. Samples were washed in PBS 1X, and fixed in 1% PFA 5 min for cells and 30min for sections. After washes (3 × 10 min in PBS 1X), samples were mounted in PBS, 20% glycerol.

### Animals

Mouse lines used in this study have been described and provided by the corresponding laboratories: C57BL/6N mice (Janvier labs) aged 8-9 week-old were used for the experiments with osmotic pumps implantations. Mice were anesthetized by isoflurane. Surgical procedures were performed under sterile conditions. Animals were handled according to national and European community guidelines, and protocols were approved by the ethics committee at the French Ministry (Project No APAFIS#26142-2020070210122646 v2).

### Muscle regeneration model, IFNγ and drug treatments

8-9 weeks old C57BL/6 male mice were obtained from Janvier labs and were maintained in a pathogen-free facility at 24 °C under a 12 h/12 h light/dark cycle with free access to food and water. The mice were weighted at day 0. Injury-induced muscle regeneration was performed by BaCl2 (0.6%) in TAs. Briefly, the mice were anesthetized by isoflurane and 50uL BaCl2 0.6% were injected in each TA (*Hardy, D. et al. PLOS ONE* ***11***, *e0147198 (2016*). Mice received IFNγ (6ug/pump; *130-105-773, Miltenyi*) or NaCl (CTL mice) by subcutaneous inserted osmotic pumps (alzet; model 1002) for 2 weeks. For drug treatment, ruxolitinib (*Selleckchem*, S1378) was dissolved in DMSO to make stock solution (60mg/mL). Ruxolitinib was diluted for oral gavage in water with 30% PEG in H2O. Each mouse received two gavage per day during 2 weeks, containing ruxolitinib (9.6 mg/kg/day) or solvent twice daily (2% DMS0, 30% PEG in H2O).

At 14 days after injury, mice were weighted and sacrificed by cervical dislocation while under anesthesia. Then, the following muscles (injured TA muscle, gastrocnemius and biceps non-injured) were dissected. Then, TAs, one gastrocnemius were mounted in tragacanth gum *(6% in water; Sigma-Aldrich)*, and frozen in isopentane precooled in liquid nitrogen. Gastrocnemius, biceps muscles were frozen in liquid nitrogen for molecular biology.

### ELISA

Blood was obtained in intracardiac. Sera were separated by centrifugation (2500 rpm, 20 min, 10°C). Each serum was assayed for murine IFN-y in duplicate using an enzyme immunoassay kit *(Peprotech, BGK01580*).

### Myofibers isolation

Single myofibers were isolated from C57BL/6N mouse EDL (*extensor digitorum longus*) muscle, as previously described protocol^30^. Muscles were dissected and digested in a filtered solution of 0.2% collagenase *(Sigma)* in DMEM (1X) + GlutaMAXTM-I *(Gibco 31966-021)* with 1% Penicillin/Streptomycin (Gibco) for 1h30 at 37°C. After connective tissue digestion, mechanical dissociation was performed to release individual myofibers that were then transferred to serum-coated Petri dishes for 20 min. Single myofibers were placed in isolation medium DMEM (1X) + GlutaMAX™-I, 20% FBS *(Foetal Bovine Serum, ref 10270 Thermo Fisher Scientific)* and 1% CEE *(Chicken Embryo Extract, CE-650-J)* with or without murine IFNγ (2×10^3^ U/mL) *(130-105-773, Miltenyi). F*our independent experiments were assayed for each data point. Then, the fibers were fixed in 4% PFA for 10 min, and washed 3 times with PBS 1X and they were stored at 4°C before immunostaining. The immunostaining was performed as described in ‘‘Immunostaining and histology” with “antibodies” described after.

### Immunostaining and histology

For immunostaining, cells were fixed in PBS, 4% paraformaldhehyde 10 min at RT, washed in PBS 1X and permeabilized with PBS 1X, 0.5% Triton X-100 10min at room temperature. After three washes in PBS 1 X, cells were blocked with Bovine Serum Albumin (BSA), 10% 30min at RT. Primary antibodies were added to cells in PBS 1X, 1% BSA for 90min at 37°C or overnight at 4°C. Cells were washed three times with PBS 1X then incubated with the secondary antibodies 1h at room temperature (RT). Before mounting with fluoromount-G *(Interchim)*, cells were washed three times with PBS 1X. For histology, the sections were kept at RT overnight before staining. Sections were then rehydrated in PBS 1X for 10 min and fixed in 4% paraformaldehyde for 10min at RT. Then, the slides were washed 2X 5 min in PBS 1X, permeabilized and blocked with 10% BSA and we added anti-mouse IgG Fab fragment (Jackson, 115-007-003) when it is necessary. Then, the slides were incubated with primary antibodies in a solution of PBS 1X, 0.1% BSA, ON at 4°C *(Table 2)*. Sections were washed in PBS 1X, 3X 5 min and incubated with the secondary antibodies 1h at RT. Sections were washed in PBS 1X, 3×5 min, and mounted with fluoromount-G *(Interchim)*.

### Clearing method for 3D myo-angiogenesis analysis

Human muscle biopsies (healthy muscle, CTL and myositis IBM/ASS) were fixed in 4% PFA for 2 hours. Then, muscles were washed three times with PBS 1X for 30 min. Microtome cutting was realized on muscle biopsies into 1 mm slices. Permeabilization was done using PBST (2% Triton X-100 in PBS) overnight at RT. Then, samples were incubated with primary antibodies anti-M-CADHERIN (R&D AF4096, 1/50), anti-DYSTROPHIN (Invitrogen PA1-21011, 1/250), and anti-MHC-II (Dako 110843-002/Clone CR3/43, 1/400) in dilution buffer (1% goat serum, 0,2% Triton-X 100, 0,2% sodium azide in PBS) for 5 days at 37°C, under agitation. Muscles were extensively washed with washing buffer (3% NaCl, 0,2% Triton X-100 in PBS). After washes, samples were incubated in secondary antibodies solution with Alexa Fluor 647-, 555-, 488-conjugated secondary antibodies (1/500, Life Technologies) for 3 days at 37°C, under agitation. Then, DAPI was added to muscle samples for 2 hours at 37°C, under agitation. After washes, they were placed in 1.52 RapiClearR reagent *(SunJiLab)* and stored at RT, out of the light.

Imaging was performed on Spinning disk microscope (Leica) using 25x water-immersion objective at Institut Jacques Monod (Paris). To control sample positioning and focus, a motorized Prior stage and piezo Z drive were used. Emission bands of 425-475 nm (blue), 500-550 nm (green), 570-620 nm (red) and 640-655 nm (far red) were used in confocal mode. Laser intensity was normalized to 80%. Image processing was performed in Fiji to normalize intensity throughout the depth and denoising. Images were then opened in Imaris 9 (Oxford Instruments) and used to do the following quantifications. Satellite cell density measurements were performed using volume plugin on Imaris.

### Antibodies for *in vitro* experiments

Primary antibodies used include: mouse anti-desmin (1/500, Dako M0760); rabbit anti-ki67 (1/250, sp6, abcam ab16667); mouse anti-Pax7 (1/100, santa-cruz sc-81648); rabbit anti-MyoD (1/200, cell signaling D8G3); rabbit anti-MyoG (1/200, santa-cruz SC576 M225); mouse anti-Myosin Heavy Chain (1/500, MF20 DSHB); Mouse anti-Human HLA-DP, DQ, DR (1 :100, Dako (110843-002/Clone CR3/43); Rabbit anti-CIITA (1/50, Thermofisher PA521031).

Alexa-conjugated secondary antibodies (1/500, Molecular Probes) and DAPI (1/5000). Following immunofluorescence, cells and sections were mounted with Fluoromount-G Mounting Media between slide and coverslip.

### Antibodies for *ex vivo* and *in vivo* experiments

Primary antibodies used include rabbit anti-ki67 (1/200, abcam sp6); mouse anti-Pax7 (1/100); Rat anti-CD68 (1/100, BD 137002 clone FA11CD31); Rat anti-MHC2 (1/300, Invitrogen 14-5321-85), mouse anti-MYH3 (1/250, Santa Cruz / sc-53091

Rabbit anti-Laminin (1/1000, Sigma L9393); Rat anti-mouse CD8 (Novus NBP1-49045SS); Rabbit anti-human CD8 (1/100, Abcam ab4055); Rat anti-CD3 (1/100, Abcam ab11089)

Alexa-conjugated secondary antibodies (1/500, Molecular Probes) and DAPI (1/5000)

Following immunofluorescence, cells and sections were mounted with Fluoromount-G Mounting Media between slide and coverslip.

### Automated morphometric analyses

Morphometric analyses were conducted using Fiji, an open-source image-processing package based on ImageJ®, in the digitized images of the entire muscle section. To automatically detect and quantify the number of myofibers and their size (diameters, CSA…) in muscle sections, we used the macro script we developed for the assessment of human and mice histological samples in collaboration with IMRB image platform (For details see ^55^).

### Western Blot

Frozen mice muscles or human muscle cells were homogenized in lysis buffer (RIPA) supplemented with β-glycerophosphate (1mM), protease inhibitor cocktail (1:100; Sigma P8340), phosphatase inhibitor cocktail (1X; *Thermo Fisher scientific, 88668*) and clarified by centrifugation. Proteins quantifications were performed by Pierce™ BCA Protein Assay *Kit (Thermo Fisher Scientific, 23225)* and an equal protein mass of 30ug in 10μL was subjected to NuPAGE™ 4-12 % Bis-Tris Midi Protein Gels *(Invitrogen™, NP0335BOX)* in Xcell4 Surelock tank *(Life Technology SAS, WR0100)* using NuPAGE™ MES SDS Running Buffer (20X) *(Invitrogen™, NP000202)*. Protein transfer to polyvinylidene difluoride (PVDF) *(Fisher Scientific, IB401032)* membrane was performed using iBlot2 Dry Blotting System *(Fisher Scientific, IB21001)* and iBlot™ 2 Transfer Stacks *(Invitrogen™, IB24001)*. Membranes were blocked in fish gelatin solution, then probed with rabbit anti-STAT1 (1:1000; 9172S, Cell signaling), rabbit anti-Phospho-STAT1 (Tyr701)(1:1000; *44376G, invitrogen)* and rabbit anti-β-tubulin (9F3) (1:1000; *#2128 Cell signaling*) overnight at 4°C. Membranes were then washed and exposed 1 hour to HRP-conjugated goat anti-rabbit (1:5000; *Santa Cruz, sc-2054*) secondary antibodies. Proteins were visualized by a chemiluminescence assay kit (*SuperSignal™ West Femto; Fisher scientific*) using a c600 scanner (*Azure Biosystems, Inc*., *Dublin, Ohio, USA*) and signals were quantified using ImageJ software (V1.52s). To perform level expression comparison, the quantification of STAT-1 was divided by the β-tubulin quantification, and Phospho-STAT1 by STAT1.

### RNA extraction and Real-Time Quantitative PCR

Total RNA was extracted from muscle sample using TRIzol and total RNA from sorted cells using a QIAGEN RNeasy Micro or Mini Kit according to the manufacturer’s instructions (QIAGEN, Hilden, Germany). RNA was quantified by Nanodrop. SuperScript III Reverse Transcriptase from the Invitrogen kit converted RNA into cDNA using the Veriti 96-Well Fast Thermal Cycler *(Applied Biosystems)*.

Gene expression was quantified by real-time qPCR with the StepOnePlus real-time PCR system *(Applied Biosystems)* using SYBR Green detection tools (Applied Biosystems). Expression of each gene was normalized to TATA Box Protein (Tbp) gene expression. Results are reported as relative gene expression (2-DDCT). Specific forward and reverse primers used in this study are listed in *Table 3*.

### RNA-sequencing of muscle

RNA was prepared as described for RNA extraction and sent to the IMRB (Institut Mondor de Recherche Biomédicale) genomic platform.

Libraries were prepared with TruSeq Stranded Total Library preparation kit (*ref 20020598*) according to supplier recommendations (Illumina, San Diego, CA). Briefly, the key stages of this protocol are successively, the removal of ribosomal RNA fraction from 500 ng of total RNA using the Ribo-Zero Gold Kit; fragmentation using divalent cations under elevated temperature to obtain ∼300 bp pieces; double strand cDNA synthesis using reverse transcriptase and random primers, and finally Illumina adapters ligation and cDNA library amplification by PCR for sequencing. Sequencing was carried out on single-end 75 bp of Illumina NextSeq500. Number of Reads 16 millions/sample.

### RNAseq analysis

The samples were quality-checked using the software FastQC (version 0.11.8). We checked that rRNA depletion had the expected quality (no prokaryotic contamination) and that more than 93% of the reads mapped to the human genome (GRCh38) using SortMeRNA (version 2.1b), FastQScreen (version 0.13) and Kraken2 (version 2.0.9 / default database). Trimmomatic (version 0.39) was used to filter reads using a quality 20 (sliding window of 5 reads) and minimal length of 50pb, which led to more than 96% of surviving reads. Filtered reads were aligned to the human genome (GRCh38) using STAR (version 2.6.1d). The mapping of the reads for the different regions of the genome and the level of gene expression was calculated using RSEM (version 1.3.2). The level of gene expression was normalized in CPM (counts per million). Differentially expressed genes were determined using edgeR package. GSEA (Gene Set Enrichment Analysis) were performed with clusterProfiler R package and GeneOntology database, specifically, Biological process ontology.

Heatmaps were created with pheatmap R package with mean normalized counts for each group.

### Statistical analysis

The data were analyzed by analysis of variance. In the event that analysis of variance justified post hoc comparisons between group means. The test for multiple comparisons is One-way ANOVA, Tukey’s multiple comparison, Two-way ANOVA Sidak’s multiple comparisons. For experiments in which only single experimental and control groups were used, group differences were examined by unpaired Student’s t test. Non-parametric, Mann-Whitney test was used. Pearson’s correlation method was used to perform correlation analysis. Data were expressed as the mean ± SEM. All statistical analyses were performed using Graph-Pad Prism *(version 6*.*0d, GraphPad Software Inc*., *San Diego, CA)*. A difference was considered to be significant at *P < 0.05, **P < 0.01, ***P < 0.001.

## Acknowledgments

This work was supported by funding from Association Française contre les Myopathies (AFM) via TRANSLAMUSCLE (PROJECT 19507). C. Hou benefited from Région Ile-de-France ARDOC Fellowship and RHU CARMMA Fellowship, and was recipient of Société Française de Myologie prize (2017). B. Periou benefited from RHU CARMMA Fellowship.

## Extended Data figure legends

Supplemental information contains three figures.

**Figure S1**

**(A)** ACP diseases

**(B)** Heatmap pathway IFN-I with the normalized reads per gene of CTL (n=5), DM (n=5), ASS (n=5) and IBM (n=4) RNA-seq data.

**(C)** Heatmap pathway IFN with the normalized reads per gene of CTL (n=5), DM (n=5), ASS (n=5) and IBM (n=4) RNA-seq data

**(D)** Gene ontology analysis for biological processes and KEGG of the downregulated (blue) genes in Inclusion body myositis (IBM) muscle compared to control (CTL) muscles. Selected enriched terms are presented according to the fold enrichment.

**(E)** Number of mTOR gene transcripts normalized in CPM (counts per million) in DIMs muscle performed by RNAsequencing.

**Figure S2**

**(A)** Representative 3D reconstruction of immunofluorescence staining of myofibers (DYSTROPHIN, green) and MHC-II (red) expressions performed on cleared thick CTL and ASS/IBM muscle sections. Z projection 60um. Champs 1062*1062 pixels.

**(B)** Representative 3D reconstruction of immunofluorescence staining of satellite cells (M-CADHERIN, yellow) and DAPI (blue) performed on cleared thick CTL and ASS/IBM muscle sections. Z projection 60um. Champs 1062*1062 pixels

**(C)** Quantification of the number of M-CADHERIN+ satellite cells per volume performed on in CTL (n=3) and ASS/IBM (n=3) muscles. Mann-Whitney U test, mean± SEM.

**Figure S3**

**(A)** Quantification of *ki67* gene expression by RT-qPCR performed on CTL and IFNγ exposed-MPCs. Mann-Whitney U test, means ± SEM

**(B)** Representative images showing scratch assay to measure cell migration, performed on CTL MPC and MPC exposed to IFNγ for 48 hours.

**(C)** Quantification of percentage of wound closure was determined between IFNγ and CTL MPCs at several times 7h, 24h and 48h.

**(D)** Luminescent Cell Viability Assay was used to determine the number of viable cells based on quantitation of the ATP level, performed on IFNγ-exposed MPCs versus CTL MPCs. One-way ANOVA, Kruskal-Wallis test.

**(E)** Quantification of membrane permeability in IFNγ-exposed cells (1-25 ng/ml) versus non-exposed cells at 24, 48, and 72 hours post-exposure, using CytoTox-Glo™ Cytotoxicity Assay. n=3 Statistics performed with One-way ANOVA, Kruskal-Wallis test (ns).

**(F)** Quantification of myogenic regulator factors *MyoG, Myf6* and *Myf5* gene expression by RT-qPCR performed on CTL and IFNγ exposed-MPCs. Mann-Whitney U test, means ± SEM

