## Supplementary figures and images for "Interferon-gamma mediates skeletal muscle lesions through JAK/STAT pathway activation in inclusion body myositis"

### Supplemental Figure 1

Supplemental Figure S1

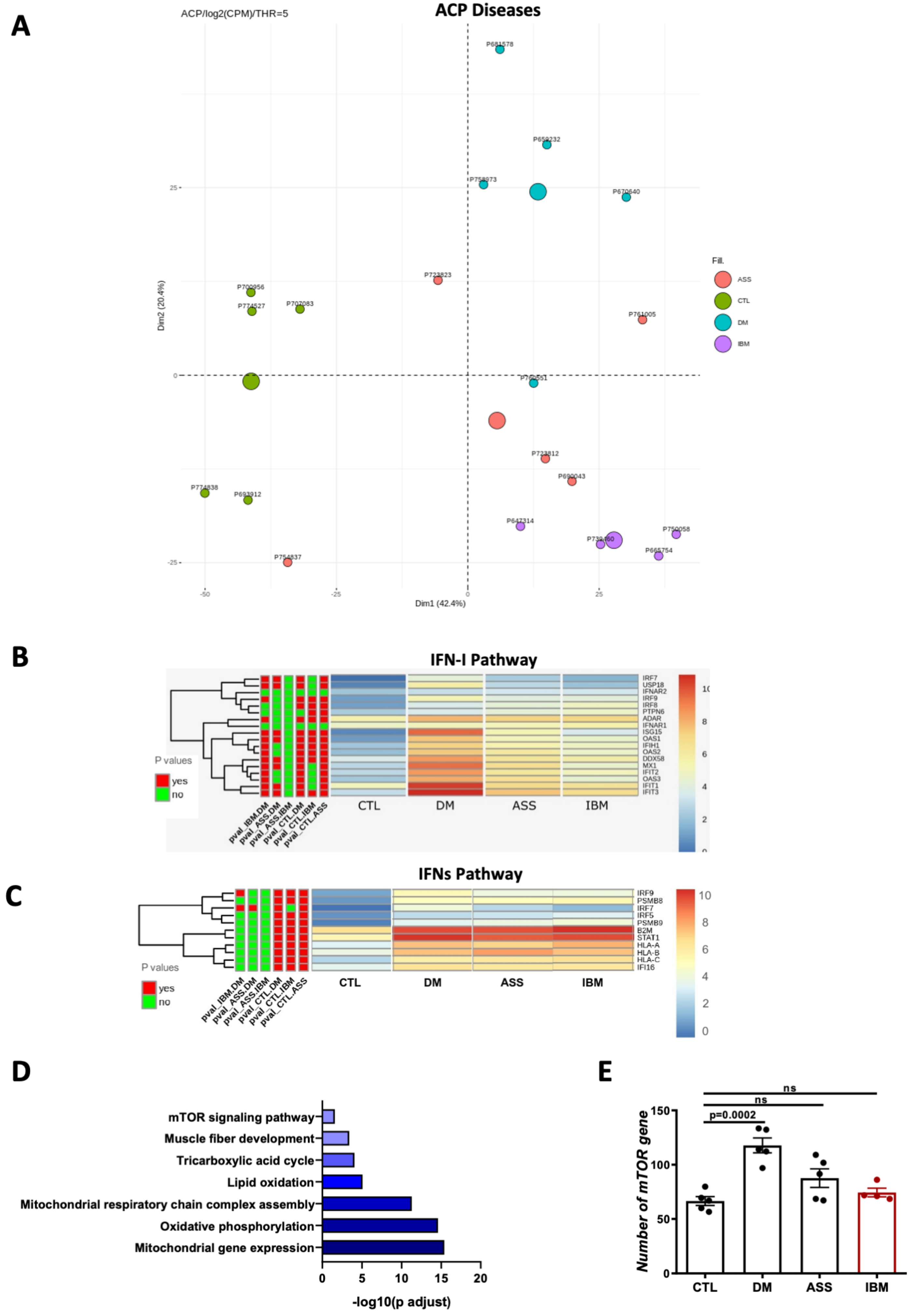

### Supplemental Figure 2

Supplemental Figure S2

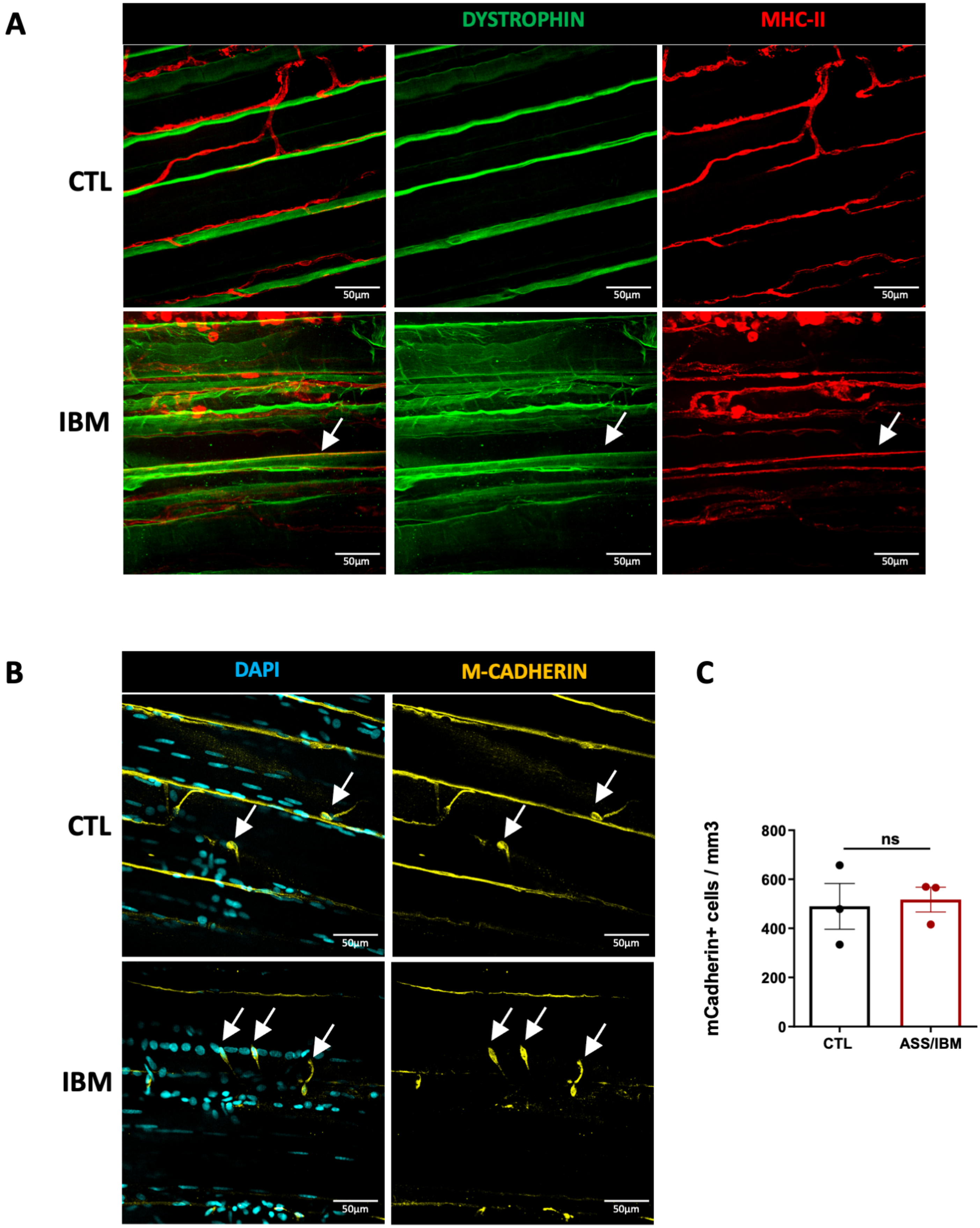

### Supplemental Figure 3

#### Supplemental figure S3 : In vitro studies

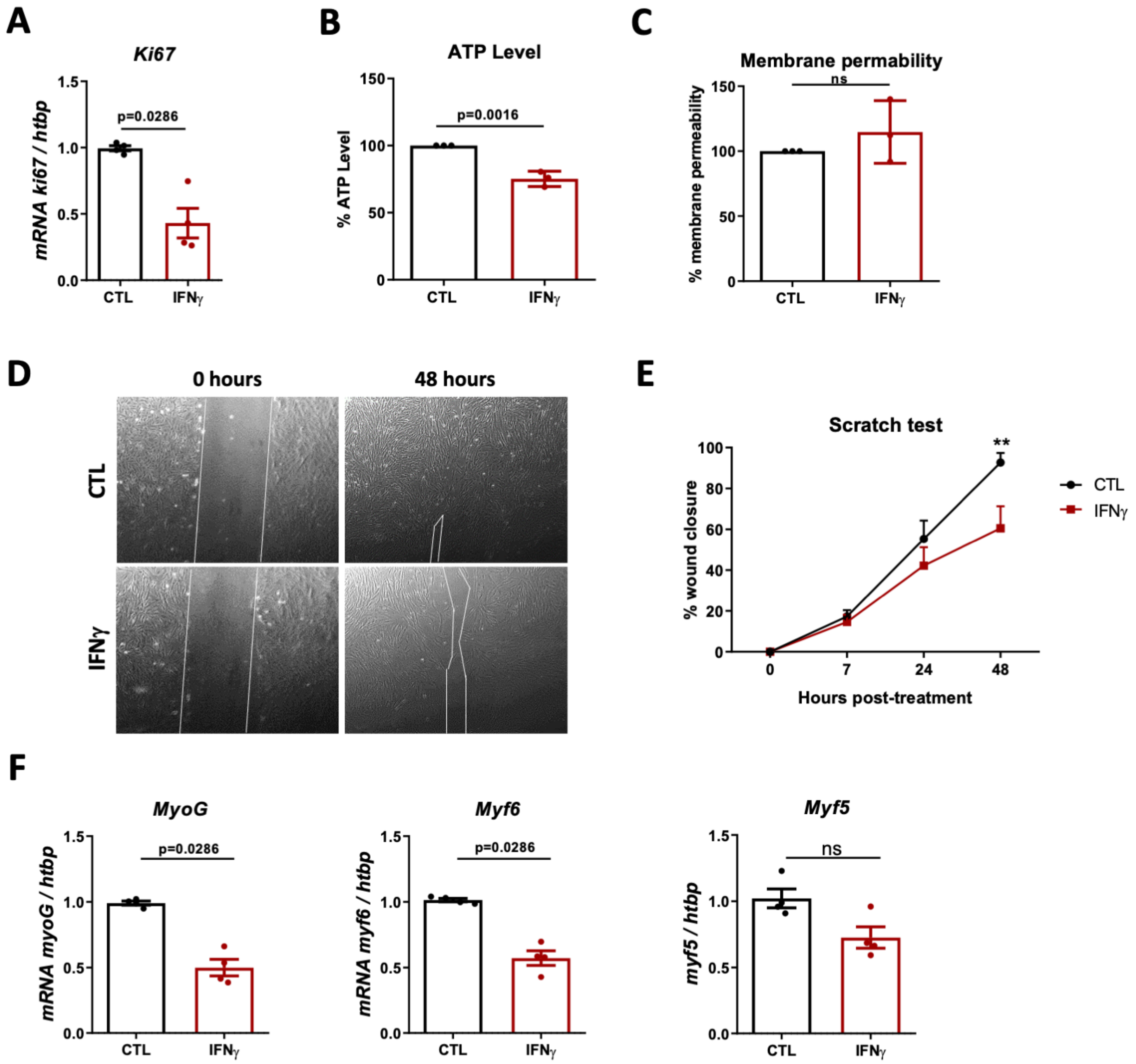
