## Supplemental Table 1 for "Interferon-gamma mediates skeletal muscle lesions through JAK/STAT pathway activation in inclusion body myositis"

| Number | Groups | Sex | Age (diagnostic) | Muscle | RNAseq | Histology |
|---|---|---|---|---|---|---|
| 1 | CTL | F | 44 | deltoid | yes | yes |
| 2 | CTL | F | 34 | deltoid | yes | yes |
| 3 | CTL | M | 51 | deltoid | yes | yes |
| 4 | CTL | F | 54 | deltoid | yes | no |
| 5 | CTL | M | 44 | deltoid | yes | no |
| 6 | CTL | M | 60 | deltoid | no | yes |
| 7 | CTL | F | 27 | deltoid | no | yes |
| 8 | CTL | M | 54 | deltoid | no | yes |
| 9 | CTL | M | 52 | deltoid | no | yes |
| 10 | CTL | M | 38 | deltoid | no | yes |
| 11 | CTL | F | 41 | deltoid | no | yes |
| 12 | CTL | M | 50 | deltoid | no | yes |
| 13 | CTL | M | 59 | deltoid | no | yes |
| 14 | CTL | M | 35 | deltoid | no | yes |
| 15 | CTL | F | 66 | deltoid | no | yes |
| 16 | CTL | M | 66 | deltoid | no | yes |
| 17 | CTL | F | 51 | deltoid | no | yes |
| 18 | CTL | F | 83 | deltoid | no | yes |
| 19 | CTL | F | 41 | deltoid | no | yes |
| 1 | DM | F | 82 | deltoid | no | yes |
| 2 | DM | F | 34 | deltoid | yes | yes |
| 3 | DM | F | 84 | deltoid | yes | yes |
| 4 | DM | F | 47 | deltoid | yes | yes |
| 5 | DM | F | 43 | deltoid | yes | no |
| 6 | DM | M | 24 | deltoid | yes | no |
| 7 | DM | F | 83 | deltoid | no | yes |
| 8 | DM | F | 48 | deltoid | no | yes |
| 9 | DM | F | 65 | deltoid | no | yes |
| 10 | DM | F | 34 | deltoid | no | yes |
| 11 | DM | M | 73 | deltoid | no | yes |
| 1 | ASS | M | 49 | deltoid | yes | yes |
| 2 | ASS | M | 41 | deltoid | yes | yes |
| 3 | ASS | F | 61 | deltoid | yes | yes |
| 4 | ASS | F | 83 | deltoid | yes | no |
| 5 | ASS | M | 59 | deltoid | yes | no |
| 6 | ASS | M | 49 | deltoid | no | yes |
| 7 | ASS | M | 55 | deltoid | no | yes |
| 8 | ASS | M | 69 | deltoid | no | yes |
| 9 | ASS | F | 71 | deltoid | no | yes |
| 10 | ASS | F | 37 | deltoid | no | yes |
| 1 | IBM | M | 68 | deltoid | yes | yes |
| 2 | IBM | M | 51 | deltoid | yes | yes |
| 3 | IBM | M | 88 | deltoid | yes | no |
| 4 | IBM | M | 69 | deltoid | yes | yes |
| 5 | IBM | M | 71 | deltoid | no | yes |
| 6 | IBM | M | 72 | deltoid | no | yes |
| 7 | IBM | M | 58 | deltoid | no | yes |
| 8 | IBM | M | 88 | deltoid | no | yes |
| 9 | IBM | F | 61 | deltoid | no | yes |
| 10 | IBM | M | 61 | deltoid | no | yes |
| 11 | IBM | M | 30 | deltoid | no | yes |
| 12 | IBM | M | 71 | deltoid | no | yes |
