## Supplemental Table 2 for "Interferon-gamma mediates skeletal muscle lesions through JAK/STAT pathway activation in inclusion body myositis"

| ANTIBODY | SOURCE / REFERENCE | IF Dilution |
|---|---|---|
| DESMIN | Dako / M0760 | 1:500 |
| KI67 | Abcam / ab16667 | 1:250 |
| PAX7 | Santa Cruz / sc-81648 | 1:100 |
| M-CADHERIN | R&D Systems / AF4096 | 1:50 |
| MYOD | Cell signaling / D8G3 | 1:200 |
| MYOG | Santa-cruz / sc576 M225 | 1:200 |
| MHC | DSHB / MF20 | 1:500 |
| MYH3 | Santa Cruz / sc-53091 | 1:250 |
| HLA-DP, DQ, DR | Dako / 110843-002 | 1 :100 |
| MHC-II | Invitrogen / 14-5321-85 | 1:300 |
| CIITA | Thermofisher / PA521031 | 1:50 |
| CD68 | BD / 137002 clone FA11CD31 | 1:100 |
| Human CD8 | Abcam / ab4055 | 1:100 |
| CD3 | Abcam / ab11089 | 1:100 |
| LAMININ | Sigma / L9393 | 1:1000 |
| DYSTROPHIN | Invitrogen / PA1-21011 | 1 :250 |
| Bodipy | Thermofisher / D3922 | 1:500 |

**Supplementary Table 1:** List of antibodies used for immunostaining.
